# Lions and brown bears colonized North America in multiple synchronous waves of dispersal across the Bering Land Bridge

**DOI:** 10.1101/2020.09.03.279117

**Authors:** Alexander T Salis, Sarah C E Bray, Michael S Y Lee, Holly Heiniger, Ross Barnett, James A Burns, Vladimir Doronichev, Daryl Fedje, Liubov Golovanova, C Richard Harington, Bryan Hockett, Pavel Kosintsev, Xulong Lai, Quentin Mackie, Sergei Vasiliev, Jacobo Weinstock, Nobuyuki Yamaguchi, Julie Meachen, Alan Cooper, Kieren J Mitchell

## Abstract

The Bering Land Bridge connecting North America and Eurasia was periodically exposed and inundated by oscillating sea levels during the Pleistocene glacial cycles. This land connection allowed the intermittent dispersal of animals, including humans, between Western Beringia (far north-east Asia) and Eastern Beringia (north-west North America), changing the faunal community composition of both continents. The Pleistocene glacial cycles also had profound impacts on temperature, precipitation, and vegetation, impacting faunal community structure and demography. While these paleoenvironmental impacts have been studied in many large herbivores from Beringia (*e.g*., bison, mammoths, horses), the Pleistocene population dynamics of the diverse guild of carnivorans present in the region are less well understood, due to their lower abundances. In this study, we analyze mitochondrial genome data from ancient brown bears (*Ursus arctos*; n = 103) and lions (*Panthera* spp.; n = 39), two megafaunal carnivorans that dispersed into North America during the Pleistocene. Our results reveal striking synchronicity in the population dynamics of Beringian lions and brown bears, with multiple waves of dispersal across the Bering Land Bridge coinciding with glacial periods of low sea levels, as well as synchronous local extinctions in Eastern Beringia during Marine Isotope Stage 3. The evolutionary histories of these two taxa underscore the crucial biogeographic role of the Bering Land Bridge in the distribution, turnover, and maintenance of megafaunal populations in North America.

## Introduction

During the Pleistocene (2.58 million to 11,700 years ago), Eastern Beringia — the area comprising Alaska and parts of Yukon Territory — was inhabited by numerous species of megafauna (1). Many of these taxa belonged to endemic New World lineages, such as the giant short-faced bear (*Arctodus simus*), Jefferson’s ground sloth (*Megalonyx jeffersonii*), and the stilt-legged horse (*Haringtonhippus francisci*) (2). However, Eastern Beringian megafaunal diversity also included non-endemic species that dispersed from Western Beringia — the area of Russia east of the Lena River — during the Pleistocene (1, 3, 4). Some of these immigrant taxa, including moose (*Alces alces*) and elk/wapiti (*Cervus canadensis*), appear to have arrived during the Last Glacial Maximum (LGM) when the Bering Land Bridge connecting Western and Eastern Beringia was most recently exposed (5–7). Other taxa apparently invaded much earlier in the Pleistocene, for example bison (*Bison* spp.) (2, 8), and mammoth (*Mammuthus* spp.) (9, 10). However, the exact timeline and processes underlying early Pleistocene dispersals are currently poorly characterized, and it remains uncertain whether the arrivals of individual species represented independent chance events or more concerted waves of species responding to changes in climate and environment.

Sea level records from the Northern Pacific indicate that the Bering Land Bridge opened and closed multiple times during the Pleistocene (11, 12), in glacial and interglacial periods respectively. During glacial Marine Isotope Stage 6 (MIS 6) around 185 thousand years ago (kya) to 135 kya, sea levels were low enough to allow the Bering Land Bridge to be uncovered (11, 13). In the subsequent MIS 5, interglacial sea levels increased to higher than present-day, flooding the Bering Land Bridge from approximately 135 to 70 kya before it re-emerged again ~70 to 60 kya during glacial MIS 4 (12). Intermittent connections may have occurred again during MIS 3, before the final emergence during MIS 2/LGM starting around 34 kya and finishing at 11 kya (12, 14).

Repeated glacial cycles also had profound effects on vegetation, which could also influence animal dispersal. For example, increased temperature during interstadials is likely to have resulted in the landscape becoming wetter, in turn facilitating the accumulation of organic matter (“paludification”) and the expansion of peatlands (15, 16). Paludification is thought to have lowered nutrient availability and favored less palatable plant species, negatively impacting megafaunal herbivore populations. Indeed, Mann *et al*. (16) observed that during interstadials in Alaska there was an initial increase in megafaunal herbivore abundance followed by a decrease coincident with peatland expansion. In addition, bone nitrogen isotopes demonstrate that the diet of horses in Alaska changed radically coincident with an increase in peatlands during Greenland Interstadial 1 (14.7-12.9 kya) (16). Changes in herbivore communities are likely to have impacted populations of megafaunal carnivores and omnivores, potentially affecting their ability to colonize or persist in Eastern Beringia through multiple glacial cycles. However, our understanding of fine-scale carnivore responses to environmental change in Eastern Beringia has been limited by their relative rarity in the fossil record. Although several studies have used ancient DNA to examine megafaunal carnivoran population dynamics (e.g., 17, 18), sample sizes have generally been small and resolution limited.

During the Late Pleistocene, a number of megafaunal carnivorans roamed Eastern Beringia, including the giant short-faced bear (*Arctodus simus*), gray wolves (*Canis lupus*), and scimitar-toothed cats (*Homotherium serum*) (1, 19). Lions (*Panthera* spp.) and brown bears (*Ursus arctos*) appear to have dispersed into northern North America from Eurasia via the Bering Land Bridge during the Pleistocene (19). Genetic data from North American lion and brown bear subfossils (preserved non-mineralized animal remains) have revealed a complicated history (17, 18, 20–22). For example, North American Pleistocene lions have been grouped into two lineages based on both fossil evidence and mitochondrial DNA, potentially representing two distinct species (or alternatively two subspecies of the extant lion): the cave lion, *Panthera (leo) spelaea*, found in both Eastern Beringia and Eurasia; and the American lion, *Panthera (leo) atrox*, found exclusively south of the North American Cordilleran and Laurentide Ice Sheets (18, 23, 24). The genetic divergence between the American lion and its relatives is estimated to have occurred ~340 kya (18), suggesting that the ancestors of the American lion invaded North America prior to MIS 6. In contrast, molecular data suggest that brown bears first colonized North America around 70 kya (around the MIS 5/MIS 4 transition), and subsequently appear to have become locally extinct in Eastern Beringia between ~35 kya and 21 kya (17, 20, 24). Genetic data from ancient lions and brown bears has so far been limited to only short fragments of mitochondrial DNA and a small number of individuals. As a result, both the timeline for invasion and the number of waves of dispersal of brown bears and lions into North America is still relatively uncertain.

To better understand the dynamics and assembly of the Eastern Beringian megafaunal carnivoran guild and their responses to climatic and environmental change, we sequenced near-complete mitochondrial genomes from 39 Pleistocene lions and 103 Pleistocene/Holocene brown bears from Eastern Beringia and Eurasia. In combination with new radiocarbon dates and previously published genetic data this allowed us to refine the phylogenetic and temporal histories of both groups and identify common drivers of dispersal and turnover.

## Results

### Brown bears

We produced 103 new near-complete (i.e., >70% coverage) mitogenomes from Pleistocene/Holocene subfossil *Ursus arctos* specimens from North America (n=53) and Eurasia (n=50), which we analyzed along with previously published sequences from 47 brown and polar bears (25–29). We used BEAST2 (30) to create a time-calibrated phylogenetic tree (Fig. 1), which was largely concordant with previous studies in grouping Beringian brown bear mitochondrial diversity into four major spatio-temporally restricted clades: clade 2 (including clade 2a, 2b, and 2c, also encompassing extant polar bears), clade 3 (including 3a, 3b, and 3c), clade 4, and clade 5 (17, 20, 22, 25, 31, 32). The temporal and geographic distributions of the different clades appear to result from dispersals into Eastern Beringia at widely different points in time.

**Figure 1.**
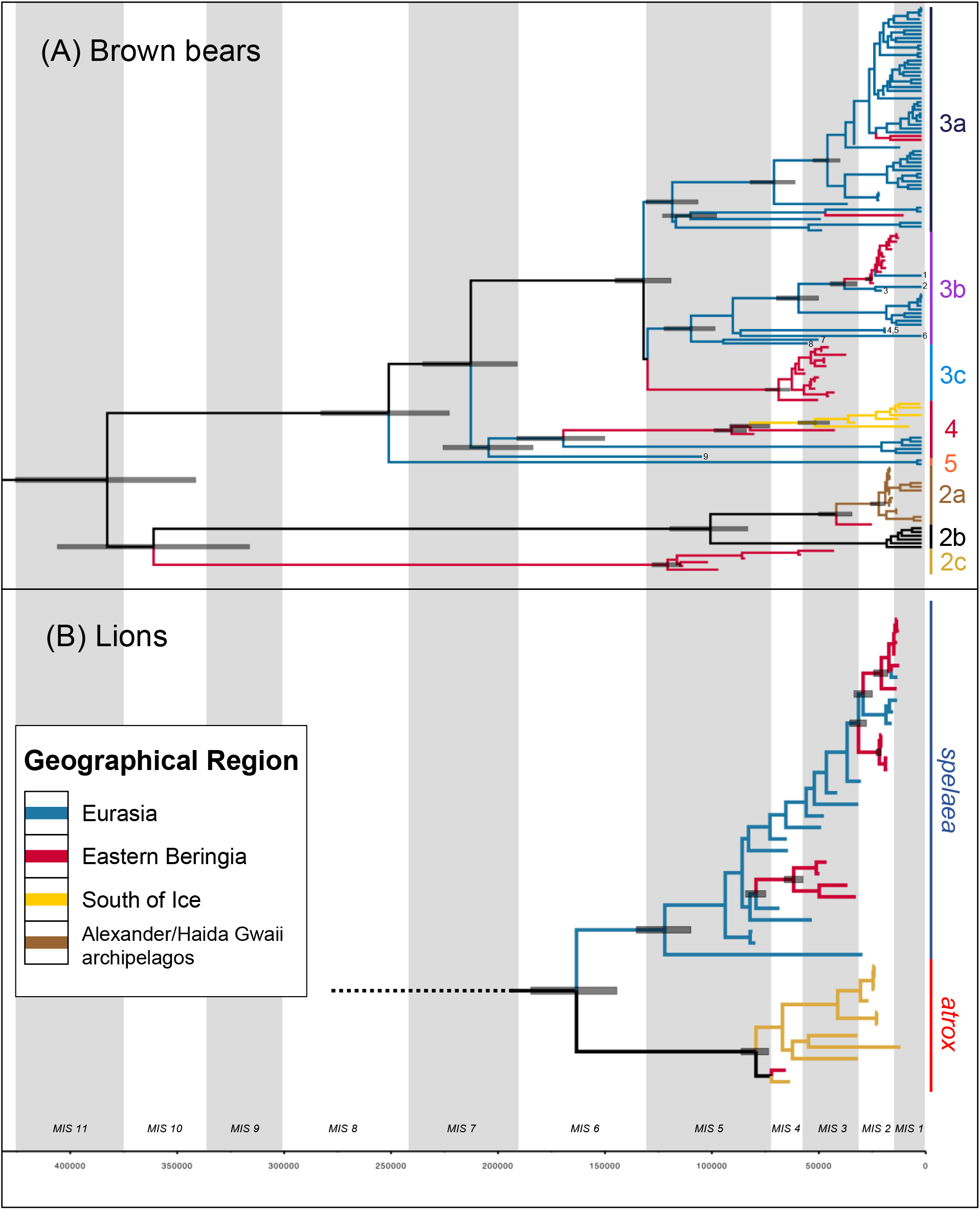
Bayesian phylogenetic trees inferred from (A) brown bear and (B) lion mitogenomes. The gray vertical columns represent odd-numbered MIS stages (interglacials) and white columns even-numbered MIS stages (glacials). Bars on nodes represent 95% Highest Posterior Densities for node age estimates indicated for modes leading to major clades and those reported in main. Numbers on tips in (A) refer to selected specimens mentioned in text: 1 = A155, 2 = A156, 3 = A1945, 4 = A1944, 5 = A1946, 6 = A138, 7 = A5889, 8 = MH255807, 9 = A5883. For detailed trees with tip labels, and posterior support values see *SI Appendix*, Fig. S4 and S5.

Within Eurasia we identified three ancient specimens (A155, A156, and A1945) with haplotypes closely related to North American clade 3b bears, and five basal Eurasian clade 3b bears (A138, A1944, A1946, A5889, and MH255807), including a published mitogenome previously assigned to clade 3c (28) (Fig. 1; *SI Appendix*, Fig. S4). The addition of these specimens increased the estimate for the Time to Most Recent Common Ancestor (TMRCA) for Eurasian and North American clade 3b bears from 75 kya (20) to 114 kya (HPD: 100.2-127.3 kya). We also identified a new haplotype that is sister-taxon to all clade 4 bears from an ancient specimen (A5883) from Da’an Cave in Northeast China, for which we estimated a median age of 103 kya (HPD: 66.7-140.6 kya).

Our time-calibrated Bayesian phylogenetic analysis returned median age estimates for five Eastern Beringian brown bear specimens that were older than the previous ~70 kya estimate for the initial colonization of North America (17, 19, 20): A345 at 78.3 kya (HPD: 58.6-98.9 kya), A335 at 82.4 kya (HPD: 64.9-103.3 kya), A298 at 95.1 kya (HPD: 64.9-127.1 kya), A193 at 100 kya (HPD: 74.0-130.2 kya), and A318 at 111.4 kya (HPD: 79.0-148.8 kya) (*SI Appendix*, Fig. S2A). These older samples likely descend from the original wave of brown bears entering North America, and all belong to either mitochondrial clade 2c or 4 (Fig. 1 & 2), neither of which is found in Eastern Beringia after 35 kya. Clade 4 bears are currently restricted to the contiguous 48 States and appear to have diverged from Eastern Beringian clade 4 bears around 83 kya (95% HPD: 73.4-93.8 kya), soon after the 92 kya TMRCA for all North American clade 4 brown bears (95% HPD: 83.2-101.6 kya). In turn, North American clade 4 brown bears appear to have diverged from Eurasian clade 4 bears (found today in Japan) much earlier, around 177 kya (HPD: 154.5-201.7 kya) during MIS 6. The other early bears, clade 2c, are currently represented by only six pre-35 kya samples from Eastern Beringia and have not been found in any modern bears, and have a TMRCA in early MIS 5, around 121 kya (HPD: 114.4-128.5 kya). An additional extinct clade, 3c, was also identified in Eastern Beringia between 40 and 35 kya, and the 15 specimens make up the majority of samples found in that time period. The TMRCA of the 15 clade 3c brown bears indicates that the clade arrived in Eastern Beringia during MIS 4 around 69 kya (95% HPD: 62.3-75.2 kya).

**Figure 2.**
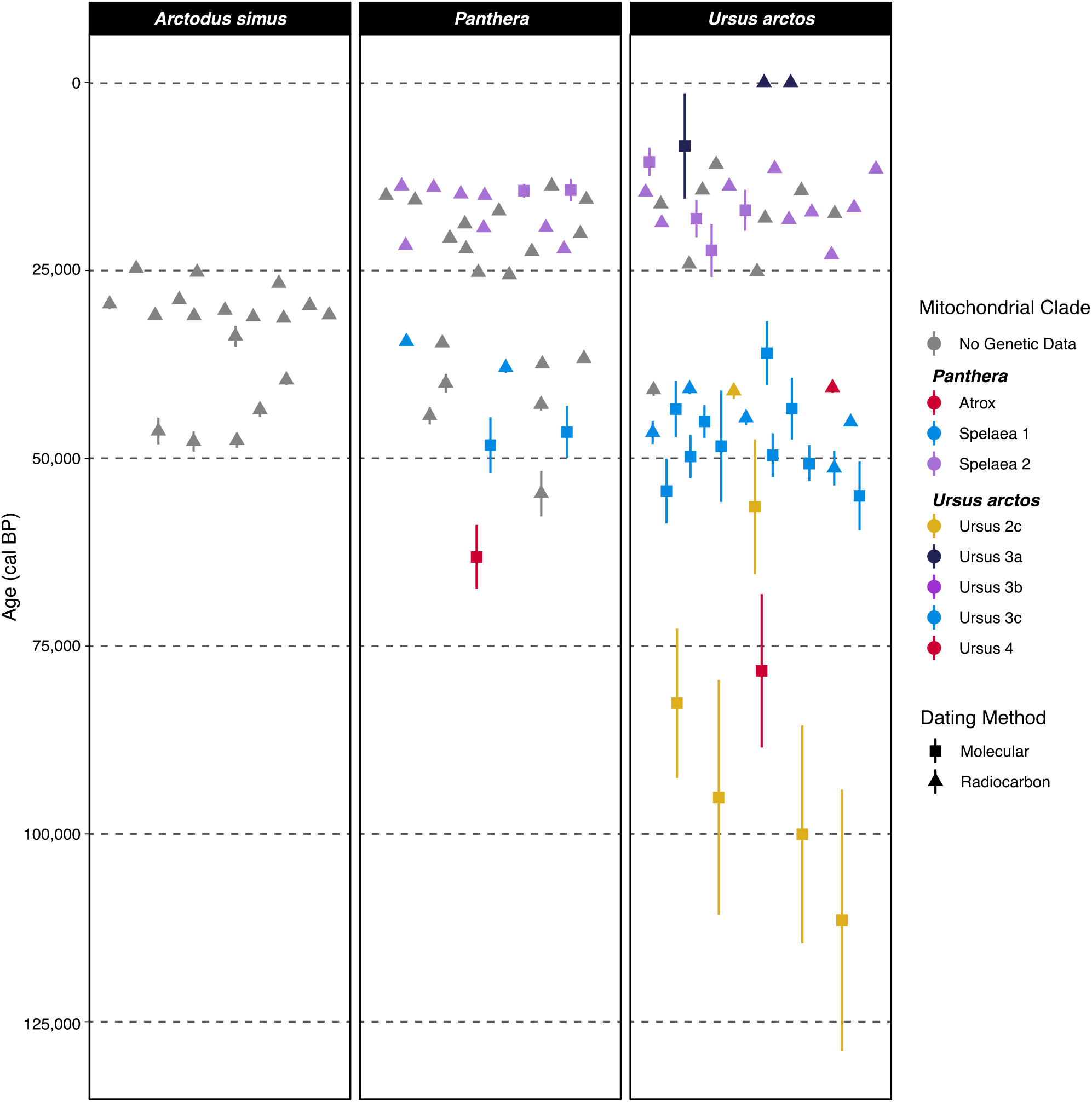
Timeline of radiocarbon and molecular dates for Eastern Beringian giant short-faced bears (*Arctodus simus*), lions (*Panthera* spp.), and brown bears (*Ursus arctos*). Dates are shown with one standard error and are colored by genetic clade. For additional radiocarbon dates used to produce this plot see *SI Appendix*, Table S1.

There is a marked gap in the Eastern Beringian fossil record of brown bears between 35 and 25 kya (Fig. 2) as previously noted (17), and after this point all samples belonged to either clade 3b or 3a. Clade 3b is the dominant group through MIS 2, comprising 13 samples, and appears to have arrived during the LGM with a TMRCA around 25 kya (95% HPD: 22.9-28.1 kya) (Fig. 1). The upper limit of this dispersal is constrained by a 39 kya estimate for the TMRCA with the closely related Eurasian clade 3b brown bears (HPD: 31.9-46.4 kya). In contrast, clade 3a is represented by only a single Holocene specimen and two previously published modern bears, and presumably constitutes a terminal-Pleistocene dispersal into North America as clade 3a bears arrive in Japan at a similar time (25).

Lastly, we recovered mitochondrial data from ten ancient clade 2a bears from Haida Gwaii and Prince of Wales Island (Alexander Archipelago). Clade 2a is closely related to the polar bear mitochondrial clade 2b, and a basal clade 2a specimen (A308) was also recovered from Engineer Creek Mine near Fairbanks, Alaska dating to 23.3 kya, the first record of clade 2a in interior Alaska. This specimen was previously reported as belonging to clade 2b using control region sequences (17, 20), although doubts about species ID (polar bear versus brown bear) and provenance have been raised (17, 33). In any case, the TMRCA of all Haida Gwaii and Alexander Archipelago specimens dates to around 20 kya (95% HPD: 17-24 kya), while the TMRCA between the Engineer Creek sample and all other clade 2a bears is 41 kya (95% HPD: 32.7-28.7 kya).

### Lions

We produced 39 new near-complete mitogenomes from lion subfossil material from North America (n=24) and Eurasia (n=15), and analyzed these along with two mitogenomes reconstructed from previously published data (34). The results of our phylogenetic analyses were in broad topological agreement with past studies, supporting the existence of two geographically restricted clades (Fig. 1) corresponding to *Panthera (leo) spelaea* (Eastern Beringia and Eurasia) and *Panthera (leo) atrox* (all other North American specimens from Edmonton southwards). We observed one important exception to this pattern: a specimen from Sixty Mile River in Yukon Territory (~64°N), A181, possessing an *atrox* (American lion) mitochondrial haplotype (Fig. 1; *SI Appendix*, Fig. S5), the first *atrox* specimen ever recorded from any locality farther north than Edmonton (~53°N). Radiocarbon dating of this specimen yielded an infinite radiocarbon age (>51,500 uncal. yBP), but our Bayesian phylogenetic analyses suggested a median age for the specimen of 67 kya (95% HPD: 51.5-84.5 kya). The TMRCA of all *atrox* lions, representing the split between the two older *atrox* specimens (>50 kya, including A181) and the younger specimens (< 35 kya), dates to MIS 5 around 81 kya (95% HPD: 74.7-87.6 kya).

Our Bayesian analysis indicated a split date between *Panthera (leo) spelaea* and *Panthera (leo) atrox* of approximately 165 kya (95% HPD: 145.0-185.2 kya). This MIS 6 divergence date is substantially younger than the previous estimate of 340 kya based on short control region sequences (18), which was likely an overestimate resulting from application of a fossil-based node-age constraint and the time-dependency of mitochondrial substitution rates (35). By relying on radiocarbon-dated tips to calibrate our analysis we have minimized the impact of rate time-dependency, allowing more accurate dating of population splits and sample ages, as demonstrated by the results of our leave-one-out cross-validation (*SI Appendix*, Fig. S1). Within Beringian lion diversity we were able to identify a genetically distinct pre-LGM mitochondrial clade of Eastern Beringian *Panthera (leo) spelaea* specimens with a TMRCA of 63 kya (95% HPD: 58.9-67.6 kya). These pre-LGM samples are genetically distinct from the two clades that include all younger Eastern Beringian lion specimens, which have TMRCAs of 23 kya (95% HPD: 22.1-24.5 kya) and 22 kya (95% HPD: 18.9-25.5 kya), and a combined TMRCA of 33 kya (95% HPD: 29.2-37.0 kya). This suggests that in addition to the original dispersal of the ancestors of *Panthera (leo) atrox*, lions appear to have dispersed into North America on at least two other occasions during the Late Pleistocene. It is notable that the hiatus in the fossil record between the pre- and post-LGM lion clades falls between 33 and 22 kya, closely mirroring the pattern of local extinction observed in brown bears (Fig. 2).

### Phylogeography: Testing the influence of the land bridge

The results of our separate phylogenetic analyses of brown bears and lions hinted at the existence of synchronous waves of dispersal and extinction tied to Pleistocene glacial cycles: in particular, most dispersal events seemed to occur during glacials, when the land bridge was present. To explicitly test whether the spatio-temporal distribution and parallel lineage turnover of lions and bears in Eastern Beringia was strongly affected by the presence or absence of the Bering Land Bridge, we performed a phylogeographic analysis in BEAST (36). To overcome low power and over-parameterization issues caused by the low number of dispersals in each clade, we used a novel approach uniting joint-tree (37), and epoch-clock (38) methods. We estimated both the bear and lion trees together in a single MCMC analysis (as separate unlinked trees); each tip in the trees (i.e., each specimen) was assigned an additional phylogeographical trait: Eurasia (Western Beringia) or North America (Eastern Beringia and South-of-the-Ice). We then estimated east-west dispersal rates (i.e., the rate of change of this phylogeographic trait) simultaneously across both the bear and lion phylogenies, along with all other parameters associated with the previous two separate analyses (i.e., clock models, substitution models, topology, branch lengths). By using a single shared biogeographic model, data from both brown bears and lions are pooled to estimate dispersal patterns and drivers (37). We compared two dispersal models using this method. (1) A simple null model, where a single dispersal rate across time was estimated, and (2) an epoch-based model where separate rates were estimated for two different groups of time slices: one rate for all periods when the Bering Land Bridge was likely emergent (i.e., glacials, even-numbered MISs) and another rate for all periods when the Bering Land Bridge was submerged (i.e., interglacials, odd-numbered MISs). Bayes factors (39) provided moderate support for the epoch-based model over the single-rate null model (BF=3.038). The estimated dispersal rate for glacials was approximately 13 times higher than the dispersal rate during interglacials (1.56E-5 versus 1.22E-6 events per lineage per year). *SI Appendix*, Fig. S6 shows the pattern driving this difference: branches containing inferred dispersals are concentrated in glacials, yet the combined glacial epochs occupy less time and shorter tree length (compared to the combined interglacials).

## Discussion

Our results demonstrate that Pleistocene glacial cycles were an important driver of population dynamics in both Eastern Beringian brown bears and lions. In particular, dispersal between Western and Eastern Beringia was heavily influenced by presence of the Bering Land Bridge, with inferred dispersal rates across both species being over an order of magnitude higher during colder periods. This result strongly implicates geographical and environmental changes caused by glacial cycles as key drivers of carnivoran diversity, which is further supported by the remarkably parallel and synchronous response to these drivers observed in both brown bears and lions. For example, the respective origins of the American lion (*atrox*) mitochondrial lineage (~165 kya) and North American clade 4 brown bear lineage (~177 kya) – the earliest representatives of both species observed in North America (Fig. 2) – occurred during MIS 6, the Illinoian glaciation (Fig. 1), when the Bering Land Bridge was likely exposed (Fig. 3A). This is consistent with the first recorded lions occurring in Sangamonian (MIS 5) deposits in Kansas and Texas (40–42). Notably, this also aligns with evidence that the steppe bison (*Bison priscus*) and red foxes arrived in North America during MIS 6 (8), or immediately prior (43, 44), respectively.

**Figure 3.**
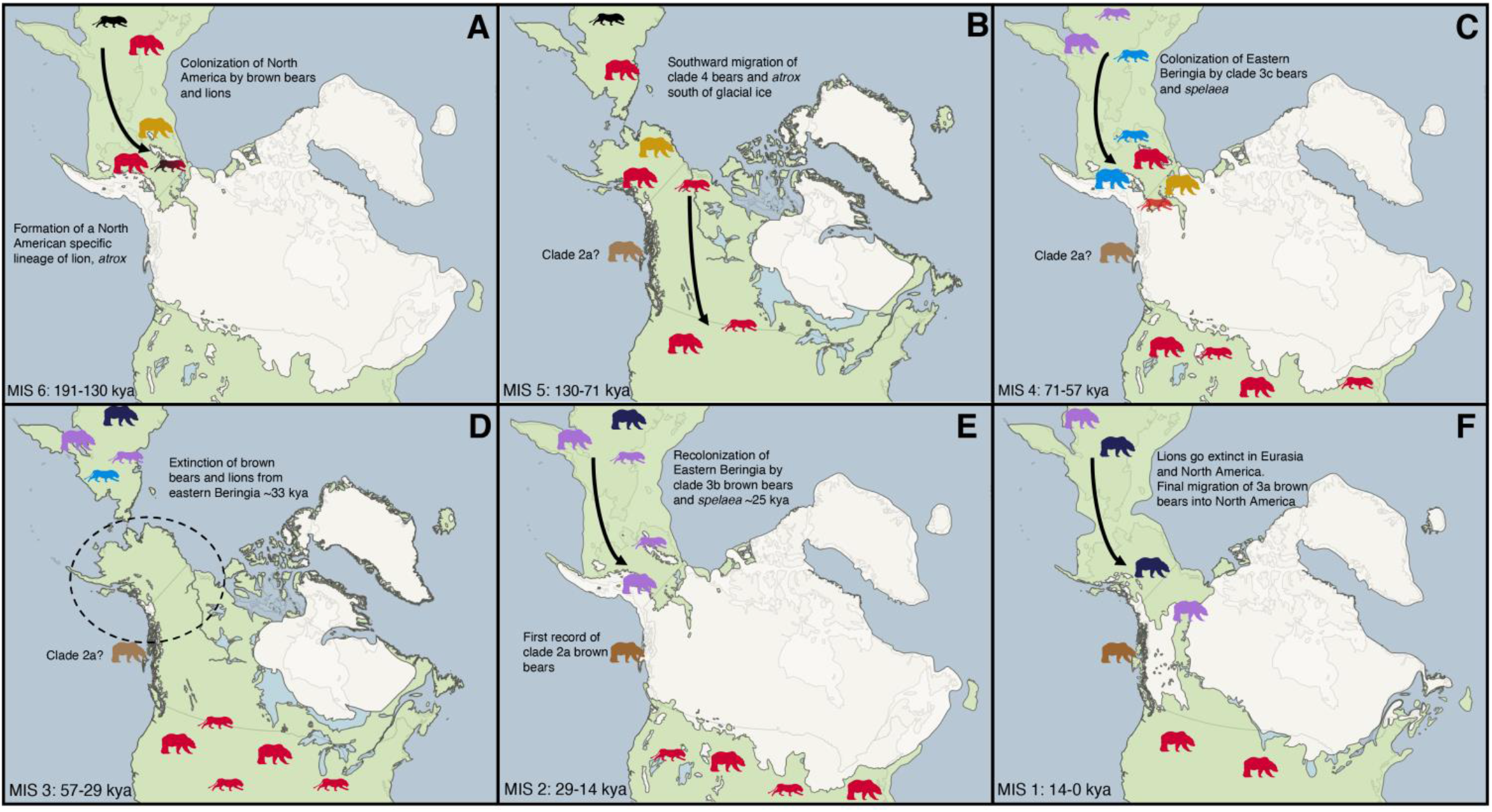
Map of Late Quaternary phylogeography of North American brown bears and lions during 6 time periods. A) MIS 6, 191-130 kya, brown bears and lions first colonize North America via the Bering Land Bridge; B) MIS 5, 130-71 kya, Bering Land Bridge is flooded, dispersal of brown bears and lions south of continental ice-sheets; C) MIS 4, 71-57 kya, dispersal of clade 3c bears and spelaea across the Bering Land Bridge; D) MIS 3, 57-29 kya, flooding of Bering Land Bridge and extinction of both carnivoran taxa in Eastern Beringia; E) MIS 2, Last Glacial Maximum, 29-14 kya, dispersal of clade 3b bears and second wave of spelaea lions; and F) MIS 1, Holocene, 14 kya to present, lions go extinct in North America and Eurasia, additionally clade 3a bears disperse into Eastern Beringia before the Bering Land Bridge is flooded for the last time. Different colored silhouettes of brown bears and lions represent different genetic clades, corresponding to clade coloring in Fig. 1 and 2. White area represents the approximate extent of glacial ice along with rough estimates of Bering Land Bridge extent during the different time periods.

While our results suggest that clade 4 bears and *atrox* lions likely arrived in Eastern Beringia around 170 kya during MIS 6, they must have dispersed southwards soon afterwards, as individuals belonging to these lineages are never observed farther north than Edmonton (~53°N) following the end of MIS 3. The TMRCAs of the North American clade 4 brown bear clade at 92 kya and *atrox* lion clade (including all North American samples) at 81 kya, both occurred during MIS 5, suggesting that both species dispersed southwards during this warmer period when ice sheets retreated and opened an ice-free north-south corridor (Fig. 3B). This movement coincides with the first southward dispersal of the bison through the ice-free corridor between late MIS 6 and early MIS 5 (2, 8, 45). The dispersal and subsequent isolation of lions south of the ice was previously thought to have initiated the divergence between the American lion (*Panthera atrox*) and cave lion (*P. spelaea*) (18). However, our discovery of a ~66.7 thousand-year-old *P. atrox* specimen north of the ice sheets in Yukon instead suggests that the formation of the endemic American lion lineage was more likely the result of their isolation in North America after the flooding of the Bering Land Bridge during MIS 5. Alternatively, this Yukon *atrox* sample could plausibly represent a migrant from south of the ice sheets, but we favor the former hypothesis as the timing of the split between *atrox* and *spelaea* coincides with the emergence of the Bering Land Bridge and there are no putative later examples of lions dispersing northwards.

Following MIS 6, the second wave of lion and brown bear dispersals into North America appears to have occurred during MIS 4 when lowered sea levels next exposed the Bering Land Bridge (Fig. 3C), corresponding with the respective TMRCAs of the North American endemic clade 3c bears and the clade comprising the four pre-LGM Eastern Beringian *spelaea* lions. However, during the interglacial period MIS 3, as the Bering Land Bridge was again submerged (12) (Fig. 3D), all lions (*atrox* and *spelaea*) and brown bears (clades 2c, 3c, and 4) appear to have become locally extinct in Eastern Beringia (Fig. 2), with *atrox* lions and clade 4 brown bears – descendants of the first wave of dispersal – surviving only in the contiguous USA and southern Canada. The absence of both brown bears and lions from the Eastern Beringian fossil record between 35 and 25 kya does not appear to be due to a taphonomic bias, as remains of the giant short-faced bear (*Arctodus simus*) are abundant during the same period (Fig. 2). Indeed, the reappearance of both lion and bear populations appears to be closely linked in time to the extinction of short-faced bears in the area, suggesting some form of competition (17, 18, 20–22). Importantly, the timing of these carnivoran extinctions in Eastern Beringia coincides with evidence for widespread vegetation change in the region, namely expansion of peatlands caused by significant paludification (15, 16, 46).

Populations of a number of megafaunal herbivores appear to have decreased during MIS 3, possibly related to the expansion of peatlands and restrictions on foraging and nutrition (16), which may have had reciprocal impacts on the megafaunal carnivores and omnivores that preyed upon them, plausibly causing the local extinction of both lions and brown bears. For example, musk-ox populations experienced a dramatic decrease in diversity and effective population size during MIS 3 (47), mammoth populations were steadily declining (48), and bison began to experience dramatic declines towards the end of MIS 3 into MIS 2 (2, 49, 50). In addition, it appears that non-caballine horses (i.e., *Haringtonhippus*) underwent a bottleneck during MIS 3 with only a single fossil specimen found in Eastern Beringia after ~31 kya (51, 52) around the time that the brown bear and lion populations went extinct. In contrast, the giant short-faced bear appears to have persisted in Eastern Beringia throughout MIS 3. It is possible that the mobility, large home range, and solitary behavior that has been proposed for the giant short-faced bear (53, 54) may have allowed them to exploit food resources that were less available to lions or brown bears.

Following MIS 3, lions and brown bears do not reappear in the fossil record of Eastern Beringia until after 27 kya, at the height of the LGM (MIS 2), when the Bering Land Bridge once again connected Eurasia and North America. This coincides with the invasion of North America from Eurasia by wapiti and moose (5, 6), and a secondary wave of bison dispersal across the Bering Land Bridge (8). The recolonizing populations were genetically distinct from those present in Eastern Beringia pre-MIS 2 as well as those south of the ice sheets, confirming that they likely comprised a new wave of dispersal from Western Beringia (Fig. 3E). This wave of megafaunal dispersals associated with the re-emergence of the Bering Land Bridge in MIS 2 may also have included early Native American human populations, who are recorded shortly afterwards in the stratigraphic record of Chiquihuite Cave in Mexico, from around 26 kya (55).

The reappearance of lions and brown bears in Eastern Beringia during MIS 2 occurred at around the same time as the local extinction of *Arctodus*, which may relate to previously proposed competition between brown bears and *Arctodus* (17, 56). The apparent timing of the extinction of *Arctodus* in Eastern Beringia around 23 kya could be linked to the sharp climatic cooling associated with Heinrich Event 2 (24.3-23.3 ka BP). Indeed, the respective TMRCAs of the newly dispersed lion lineages at 23 kya and 22 kya suggest that they may also have been impacted by Heinrich Event 2, perhaps through population bottlenecks removing previous genetic diversity. In any case, the fact that Eastern Beringia was not instead recolonized by *atrox* lions and clade 4 bears from the contiguous USA may either reflect that conditions had not improved sufficiently to support lion and brown bear populations in Eastern Beringia before the ice-free corridor closed during the LGM or suggest that some other geographical or biogeographical barrier prevented dispersal from south of the ice sheets. Concordantly, in bison there is little evidence for northward dispersal through the ice-free corridor until after the LGM when a pulse of south to north dispersal is observed (45).

All modern and ancient clade 2a brown bears from the Alexander and Haida Gwaii archipelagos coalesce at 20 kya (95% HPD: 17.0-24.0 kya), comparable to the TMRCAs for Beringian clade 3b bears and *spelaea* lions. This supports the model proposed by Cahill *et al*. (57) for the origin of clade 2a bears, under which the mitochondrial lineage was captured by brown bears following male-biased gene-flow into a population of polar bears stranded in the Alexander archipelago after the retraction of ice sheets post-LGM. Assuming all our ancient Alexander and Haida Gwaii archipelago samples represent brown bears (or at least brown-polar hybrids), and that mtDNA diversity in the stranded polar bear population was low, the coalescence of our samples can be considered a proxy for the minimum age of hybridization between polar and brown bears, and hence a minimum age for the arrival of brown bears in the Alexander and Haida Gwaii archipelagos post-LGM. If this is the case, then brown bears arrived in the islands no later than 17 kya (the lower bound of the 95% HPD). That timing is coincident with the first records of brown bears on the Haida Gwaii archipelago ~ 17.5 kya (58) and the existence of unglaciated western Alaskan coastline, which represents an alternative southward dispersal pathway into the continent that may also have been exploited by humans (59, 60).

Overall, our results highlight the key role of Pleistocene glacial cycles in driving the distribution and diversity of North American carnivorans. Glacial cycles may also have driven parallel waves of dispersal in other regions, such as across the Sakhalin land bridge that connected Japan with mainland Asia. Genetic evidence from modern Japanese brown bears suggests multiple waves of Pleistocene dispersal in a similar temporally staggered sequence, with present day Japanese mitochondrial diversity closely mirroring that observed in modern Eastern Beringia (i.e., clades 3a, 3b, and 4) and also exhibiting a marked phylogeographic structure (25). Analysis of ancient Japanese brown bear specimens might allow determination of whether extinct Eastern Beringian clades such as 3c were also present in Japan during the late Pleistocene.

## Conclusion

Lions and brown bears display remarkably synchronous responses to Pleistocene glacial cycles, and combining phylogenetic data from these two Pleistocene carnivoran species in a shared common biogeographic model provides power to demonstrate a 13-fold increase in dispersal rate between Eastern and Western Beringia when the land bridge is present. By combining additional ancient DNA datasets from other species with trans-Beringian Pleistocene distributions (e.g., foxes), future studies may further refine the timing and magnitude of waves of dispersal across the Bering Land Bridge. A similar combined biogeographical approach may also be useful for exploring the timing of faunal dispersals through the ice-free corridor between the North American ice sheets, which available data suggests are biased southwards, with few observed northward dispersals. However, this apparent bias may be due to many ancient DNA studies focusing on recently immigrated taxa (e.g., brown bears, bison, wapiti, humans) for which Eastern Beringia acts as a source, with the contiguous USA likely a sink. Endemic North American species may exhibit different patterns of phylogeography and dispersal, and large ancient DNA datasets from species like the giant short-faced bear or the western camel (*Camelops hesternus*) would be valuable in evaluating this possibility. Our densely-sampled study of rarer carnivorans contributes to the growing body of research suggesting remarkably concerted responses to Pleistocene geographical and environmental changes across many megafaunal taxa (e.g., 61).

## Materials and Methods

### Sample preparation, DNA extraction, library preparation, and mitochondrial enrichment

We sampled 120 brown bear subfossil bone and tooth specimens from northern Asia and North America, and 47 lion subfossils from Europe, northern Asia, and North America (see Datasets S1 and S2). Twenty-six samples were radiocarbon dated at the Oxford Radiocarbon Accelerator Unit of the University of Oxford. All radiocarbon dates were calibrated with the IntCal13 curve (62) using OxCal 4.4 (63).

Sample preparation, DNA extraction and library construction were conducted in purpose-built aDNA clean-room facilities at the University of Adelaide’s Australian Centre for Ancient DNA (ACAD) or the Henry Wellcome Ancient Biomolecules Centre at the University of Oxford and a number of precautions taken to minimize contamination of samples with exogenous DNA (64). DNA extraction was performed on bone or tooth powder using either an in-house silica-based extraction protocol adapted from Dabney *et al*. (65) or a phenol-chloroform-based extraction protocol from Bray *et al*. (66). Double-stranded Illumina libraries were constructed following the protocol of Meyer *et al*. (67) with truncated Illumina adapters with unique dual 7-mer internal barcodes added to allow identification and exclusion of any downstream contamination. Partial uracil-DNA glycosylase (UDG) treatment (68) was included to restrict cytosine deamination to terminal nucleotides. Brown bear libraries were enriched with home-made RNA baits following Richards *et al*. (69) produced from long-range PCR fragments amplified from modern brown bear DNA using primers from Hwang *et al*. (70). For lion libraries, commercially synthesized biotinylated 80-mer RNA baits (Arbor Biosciences, MI, USA) were used to enrich for mammalian mitochondrial DNA (71). DNA-RNA hybridization enrichment was performed according to manufacturer’s recommendations (MYbaits protocol v3). Libraries were pooled and sequenced on an Illumina NextSeq using 2 x 75 bp PE (150 cycle) High Output chemistry. A more detailed description of the laboratory methods is available in the *SI Appendix*.

### Data processing

Sequenced reads were demultiplexed using SABRE (https://github.com/najoshi/sabre) and were then processed through Paleomix v1.2.12 (72), with adapter sequences removed and pair end sequences merged using ADAPTER REMOVAL v2.1.7 (73), and merged reads mapped against either the mitochondrial genome of Panthera spelaea (KX258452) or Ursus arctos (EU497665) using BWA v0.7.15 (74). Reads with mapping Phred scores less than 25 were removed using SAMTOOLS 1.5 (75) and PCR duplicates were removed using “paleomix rmdup_collapsed” and MARKDUPLICATES from the Picard package (http://broadinstitute.github.io/picard/). Data from our lion samples exhibited signals consistent with the presence of nuclear mitochondrial DNA segments (numts), which are known to be widespread in felid genomes (76). The numt sequence was identified and lion samples were remapped with the numt sequence included as an additional scaffold to allow separation of true mitochondrial sequences and numt sequences. Mapped reads were visualised in in Geneious Prime v2019.0.4 (https://www.geneious.com) and we created a 75% majority consensus sequence, calling N at sites with less than 3x coverage. Published sequencing data from one modern brown bear (29) and two ancient cave lions (34) were also processed through the pipeline described above (*SI Appendix*, Table S2). A more detailed description of the data processing methods is available in the *SI Appendix*.

### Phylogenetic analyses

Brown bear consensus sequences were aligned using MUSCLE v3.8.425 (72) in Geneious Prime v2019.0.4 with an additional 46 brown bear and polar bear mitogenomes downloaded from GenBank (*SI Appendix*, Table S3). Lion sequences were aligned separately as above. PartitionFinder 2.1.1 (73) was used to find the best-fitting partitioning scheme using the Bayesian information criterion, separating the data into 5 partitions for each alignment (*SI Appendix*, Table S4). Bayesian tip-dating analyses were then performed on each taxon using BEAST 2.6.1 (30). The temporal signal in our dataset was evaluated using leave-one-out cross-validation (e.g. 74), using only the finite-dated specimens. The ages of undated specimens were then estimated one at a time using the dated specimens as calibration for the molecular clock. Once all samples were assigned an age (either based on radiocarbon dating or Bayesian date estimation), we conducted a date-randomization test (74, 75). Runs described above were performed with a strict clock with a uniform prior on rate (0-10-5 mutations per site per year), constant population coalescent tree prior with a 1/x distribution on population size, a uniform prior (0-500,000) on the age of the sequence being estimated if required, and run for 30 million steps with sampling every 3000 steps. Convergence was checked in Tracer v1.7.1 (76). Final BEAST analyses were conducted using a strict clock with a uniform prior on rate (0-10-5 mutations per site per year), and a Bayesian skyline coalescent tree prior. We ran three independent MCMC chains, each run for 50 million steps, sampling every 5,000 steps. Results from individual runs were combined using LogCombiner after discarding the first 10% of steps as burn-in. Maximum clade credibility trees were generated in TreeAnnotator using the median node age.

To test for the association of migrations between Eurasia and North America with glacial periods, phylogeographic model testing was performed in BEAST (36). The same substitution model settings were used as described above, but the alignments were combined in a single analysis (excluding clade 2 bears), with a separate tree estimated simultaneously for each taxon. Each tip was assigned a binary phylogeographic character (Eurasia vs North America), and the rate of evolution of this character was estimated directly from the data. Two models for the evolution of this character were tested: a strict clock, where rates of evolution were constant through time and a two-epoch clock that had two separate rates (interglacial and glacial periods). Bayes Factors were estimated and compared using AICM (Tracer). Four independent MCMC chains were run for 20 million steps each, sampling every 2,000 steps. We checked for convergence and sufficient sampling of parameters in Tracer v1.7.1 (76). A more detailed description of the phylogenetic analysis methods is available in the *SI Appendix*.

## Acknowledgements

We would like to thank the following institutions for allowing access to specimens in their collections: University of Alaska Fairbanks Museum, University of Kansas Natural History Museum, University of Wyoming Geological Museum, Yukon Government, American Museum of Natural History, Cincinnati Museum, Bureau of Land Management Nevada-Elko District, St. Petersburg Institute of Zoology, Krakow Institute of Zoology, the Russian Academy of Sciences, Palaeontological Institute Moscow, Zoological Museum of Moscow University, The Institute of Plant and Animal Ecology of the Ural Branch of the Russian Academy of Sciences, Natural History Museum Stuttgart, University of Vienna, Museum of Natural History Vienna, Idaho Museum of Natural History, Royal Alberta Museum, Parks Canada, the Canadian Museum of Nature, Gwaii Haanas National Park Reserve and the Haida Nation. In addition, we are grateful to the following individuals who helped to collect and identify specimens and/or provided laboratory support during the early stages of the project: L. Orlando, T. Heaton, K. Chen, I. Barnes, A. Derevianko, E. Pankeyeva, I. Chernikov, M. Shunkov, A. Sher, N. Ovodov, C. Beard, D. Miao, D. Burnham, L. Vietti, M. Clementz, G. Zazula, P. Matheus, P. Wrinn, D. McLaren, and J. Austin. Gaadu Din Haida Gwaii fieldwork was funded by Social Science and Humanities Research Council of Canada Standard Grant awarded to DF (410-2005-0778). This research was funded by an Australian Research Council Laureate Fellowship awarded to AC (FL140100260) and U.S. National Science Foundation grant (EAR/SGP# 1425059) awarded to JM and AC.

