## Supplementary Information Appendix for "Lions and brown bears colonized North America in multiple synchronous waves of dispersal across the Bering Land Bridge"

**Extended Material and Methods**

**Sampling:**

We sampled 120 brown bear subfossil bone and tooth specimens from northern Asia and North America, and 47 lion subfossils from Europe, northern Asia, and North America (see Supplementary Tables 1 and 2). Fourteen brown bears specimens, and 12 lion specimens were radiocarbon dated at the Oxford Radiocarbon Accelerator Unit of the University of Oxford. All radiocarbon dates were calibrated with the IntCal13 curve (1) using OxCal 4.4 (2).

All pre-PCR steps (extraction, library preparation) were conducted in purpose-built aDNA clean-room facilities at the University of Adelaide's Australian Centre for Ancient DNA (ACAD) or the Henry Wellcome Ancient Biomolecules Centre at the University of Oxford, spatially separated and physically isolated from any other molecular laboratories. Strict protocols were followed and a number of precautions taken to minimize contamination of samples with exogenous DNA (1). Protective clothing was worn, including: hooded coveralls over ancient-DNA lab-dedicated clothing (clothes never previously worn in any other molecular laboratory), hairnets, facemasks, face shields, designated footwear for both transitional areas and the physical laboratory, and three pairs of gloves worn at all times to prevent skin exposure between frequent changes of the outer layer of gloves. Furthermore, the lab was designed with positive air pressure, flowing from the cleanest workrooms to the outside of the lab. Stringent decontamination procedures were also adhered to, including cleaning equipment and surfaces with bleach or disinfectant detergent before and after use as well as regular UV irradiation of surfaces. These precautions also included negative controls for both DNA extraction and PCR setup. PCR amplification and all downstream procedures (e.g., quantification and hybridization enrichment) were carried out in independent DNA laboratories.

**DNA Extraction:**

Potential surface contamination on each sample was reduced by UV irradiation for 15 min each side, followed by abrasion of the exterior surface (c. 1 mm) using a Dremel tool and a disposable carborundum disk. The sample was then pulverized using either a metallic mallet or a mikro-dismembrator (Sartorius). Approximately 100 mg of powder was extracted using one of two protocols: 1) Phenol-chloroform-based extraction protocol from Bray et al. (2); or 2) an in-house silica-based extraction protocol adapted from Dabney et al. (3). For the latter protocol, the powder was digested first in 1 mL 0.5 M EDTA for 60 min, followed by an overnight incubation in 970 μL fresh 0.5 M EDTA and 30 μL proteinase K (20 mg/ml) at 55°C. The samples were centrifuged and the supernatant mixed with 13 mL of a modified PB buffer (12.6 mL PB buffer (Qiagen), 6.5 μL Tween-20, and 390 μL of 3M Sodium Acetate) and bound to silicon dioxide particles, which were then washed two times with 80% ethanol. The DNA was eluted from silica particles with 100 μL TE buffer.

**Library Preparation:**

Double-stranded Illumina libraries were constructed following the protocol of Meyer et al. (4) from 25 μL of DNA extract, with truncated Illumina adapters with unique dual 7-mer internal barcodes added to allow identification and exclusion of any downstream contamination. In addition, all samples underwent partial uracil-DNA glycosylase (UDG) treatment (5) to restrict cytosine deamination, characteristic of ancient DNA, to terminal nucleotides, while eliminating damage in the centre of the molecules. A short round of PCR using PCR primers complementary to the adapter sequences was performed to increase the total amount of DNA. Cycle number was determined via rtPCR and each library split into 8 separate PCR reactions to minimize PCR bias and maintain library complexity. Each PCR of 25 μL contained 1× HiFi buffer, 2.5 mM $MgSO_4$, 1 mM dNTPs, 0.5 mM each primer, 0.1 U Platinum Taq Hi-Fi polymerase and 3 μL DNA. The cycling conditions were 94 °C for 6 min, 9-31 cycles of 94 °C for 30 s, 60 °C for 30 s, and 68 °C for 40 s, followed by 68 °C for 10 min. PCR replicates were pooled and products were then purified using AxyPrep™ magnetic beads (Axygen™). DNA was eluted in 30 μL EB buffer and quantified with a Qubit fluorometer (Thermo Fisher).

**Mitochondrial Enrichment:**

For lion libraries, commercially synthesized biotinylated 80-mer RNA baits (Arbor Biosciences, MI, USA) were used to enrich for mammalian mitochondrial DNA (6). DNA-RNA hybridization enrichment was performed according to manufacturer's recommendations (MYbaits protocol v3) with the exception that 1.25 μL of baits per reaction was used and the incubation step which was changed to 55 °C for 15 hr followed by 50 °C for 16 hrs. The beads were washed three times with 0.1 x SSC and 0.1% SDS (5 min 55 °C).

Brown bear libraries were enriched with home-made RNA baits following Richards et al. (7). DNA was extracted from two brown bear tissue samples obtained from the University of Alaska Fairbanks Museum using a Qiagen DNeasy Blood and Tissue kit following manufacturer's protocols. The mitochondrial genomes were then amplified from the two specimens in two long-range PCR fragments of 8-9 kb fragments using primers adapted from Hwang et al. (8), ensuring a T7 promoter sequence was ligated to the 5' end of one primer of each pair. The primer sequences were as follows: fragment 1, S-LA-16S-L-T7: 5'-AATTGTAATACGACTCACTATAGGG GAT GTT GGA TCA GGA CAT CCT AAT GGT GCA-3', H-12193-Leu: 5'-AGT TGC ACC AAT TTT TTG GTT CCT AAG ACC-3', and fragment 2, L-12193-Leu-T7: 5'-AATTGTAATACGACTCACTATAGGG GGT CTT AGG AAC CAA AAA ATT GGT GCA ACT-3', S-LA-16S-H, 5'-TGC ACC ATT AGG ATG TCC TGA TCC AAC ATC-3'. The long-range PCR fragments from both samples were then pooled in equimolar amount and subjected to *in vitro* transcription. The resulting RNA was then fragmented and biotinylated to form completed RNA baits specific to the brown bear mitochondrial genome. Brown bear samples were then enriched for mitochondrial DNA following the same protocol as per the lions but using the homemade RNA baits instead of commercially synthesized baits.

Full-length Illumina sequencing adapters were then added to the enriched libraries via a final round of "off-bead" PCR split into 5 replicate PCRs (25 μL) containing 1× Gold PCR buffer, 2.5 mM $MgCl_2$, 1 mM dNTPs, 0.5 mM each primer and 0.1 U AmpliTaq Gold. Cycling conditions were as follows: 94 °C for 6 min; 15 cycles of 94 °C for 30 s, 60 °C for

30 s, 72 °C for 45 s; and 72 °C for 10 min. Following PCR, replicates were pooled and purified using AxyPrep™ magnetic beads, eluted in 30 µL $H_2O$ quantified on TapeStation (Agilent Technologies). Libraries were pooled and sequenced on an Illumina NextSeq using 2 x 75 bp PE (150 cycle) High Output chemistry.

**Data processing:**

Sequenced reads were demultiplexed using SABRE (https://github.com/najoshi/sabre) using the unique 5' and 3' barcodes allowing one mismatch in the barcode sequence (-m 1). Demultiplexed reads were then processed through Paleomix v1.2.12 (9). Within Paleomix, adapter sequences were removed and paired end reads merged using ADAPTER REMOVAL v2.1.7 (10), trimming low-quality bases (<Phred20 --minquality 4) and discarding merged reads shorter than 25 bp (--minlength 25). Read quality was visualized before and after adapter trimming using fastQC v0.11.5 (http://www.bioinformatics.babraham.ac.uk/projects/fastqc/) to ensure efficient adapter removal. Merged reads were mapped against the mitochondrial genome of *Panthera spelaea* (KX258452) and *Ursus arctos* (EU497665) using BWA v0.7.15 (11) (aln -l 1024 (seed inactivated), -n 0.01, -o 2). Reads with mapping Phred scores less than 25 were removed using SAMTOOLS 1.5 (12) and PCR duplicates were removed using "paleomix rmdup_collapsed" and MARKDUPLICATES from the Picard package (http://broadinstitute.github.io/picard/).

Heterozygous sites were observed across a 7-kb region in multiple *Panthera* samples, presumably as the result of nuclear mitochondrial DNA segments (numts), which are known to be widespread in felids (13). To counteract this, we constructed a numt sequence reference by identifying runs of sequences that disagreed with flanking homozygous sequences of the "true" mitochondrial genome. This numt reference was included as an additional scaffold when mapping to the lion mitochondrial genome reference so that reads preferentially mapping to the numt reference could subsequently be excluded from downstream analyses.

Following mapping, reads for all samples were visualized in Geneious Prime v2019.0.4 (https://www.geneious.com) and we created a 75% majority consensus sequence, calling N at sites with less than 3x coverage. We also re-analyzed published data from one modern brown bear (14) and two ancient cave lions (15) through the pipeline described above to produce full mitochondrial genomes (Supplementary Table 4).

**Phylogenetic analysis:**

Using MUSCLE v3.8.425 (16) in Geneious Prime v2019.0.4, we aligned the 104 brown bear consensus sequences described above with an additional 46 brown bear and polar bear mitogenomes downloaded from GenBank (Supplementary Table 5). We aligned our lion consensus sequences the same way, thus creating a separate alignment for each taxon (i.e., *Panthera* and *Ursus arctos*)

Bayesian tip-dating analyses were performed using BEAST 2.6.1 (17) on each alignment to co-estimate the tree topology and divergence dates of our sequences. First, PartitionFinder 2.1.1 (18) was used to find the best-fitting partitioning scheme using the Bayesian information criterion, separating the data into 5 partitions for each alignment (Supplementary Table 6). We then evaluated the temporal signal in our dataset using leave-one-out cross-validation (e.g. 19), using only the finite-dated specimens (28 lions and 119 brown bears). In sequential analyses we left out and then attempted to estimate the age of each specimen. For all but two of the lion and two of the brown bear specimens, the “true” (radiocarbon) age was within the 95% credibility interval of the estimated age, suggesting that our dataset included sufficient temporal information to estimate the age of undated samples (Supplementary Figure 1). Consequently, we performed sequential analyses where undated samples were added to the dataset one at a time, in order to estimate their ages (Supplementary Figure 2). Runs were performed with a strict clock with a uniform prior on rate (0-$10^{-5}$ mutations per site per year), constant population coalescent tree prior with a 1/x distribution on population size, a uniform prior (0-500,000) on the age of the sequence being estimated, and run for 30 million steps with sampling every 3000 steps. Some chains were extended to ensure effective sampling sizes near or above 200 for all parameters. The first 10% of samples were discarded as burn-in and parameter values were monitored to check for convergence in Tracer v1.7.1 (20). Once all samples were assigned an age (either based on radiocarbon dating or Bayesian date estimation), we conducted a date-randomization test (19, 21). Runs were conducted as for date estimation but excluding a prior on sequence age. For both datasets the rate estimate of the original data did not overlap the credibility intervals of the rate estimate from 20 randomized replicates (Supplementary Figure 3), suggesting that our dataset could be used to reliably estimate evolutionary rate and divergence times.

For the final BEAST analysis, a strict clock was used with a uniform prior on rate ($0\text{-}10^{-5}$ mutations per site per year), and a Bayesian skyline coalescent tree prior. We ran three independent MCMC chains, each run for 50 million steps, sampling every 5,000 steps. We checked for convergence and sufficient sampling of parameters in Tracer v1.7.1 (20) and combined individual runs in LogCombiner, after discarding the first 10% of steps as burn-in. MCC consensus trees were generated in TreeAnnotator using the median node age.

**Phylogeographic model testing:**

The joint-tree epoch analyses used BEAST (22) and identical DNA substitution model settings as above. However, the analysis was set up so that two separate alignments (lions and bears) were encoded in a single common xml file, and two separate trees (lions and bears) were estimated simultaneously during the MCMC. Additionally, clade 2 bears were excluded from the analysis due to a lack of sufficient sampling and introgressed relationship with polar bears (*Ursus maritimus*). Each tip or taxon was coded with an additional binary phylogeographic character (Eastern vs Western Beringia), and the rate of evolution of this character was estimated directly from the data. Two models for the evolution of this character were tested: a strict clock, where rates of evolution were constant through time, and a two-epoch clock, which had two separate rates (interglacial and glacial periods). We ascertained Bayes Factors using both stepping-stone and AICM (Tracer) approaches, but the former values were very unstable across runs (possibly due to poor convergence during some steps), and the values reported in the main text are AICM.

We ran four independent MCMC chains, each run for 20 million steps, sampling every 2,000 steps. We checked for convergence and sufficient sampling of parameters in Tracer v1.7.1 (20) and combined individual runs using LogCombiner after discarding the first 20% of steps as burn-in. MCC consensus trees were generated in TreeAnnotator using the median node age (Supplementary Figure 6).

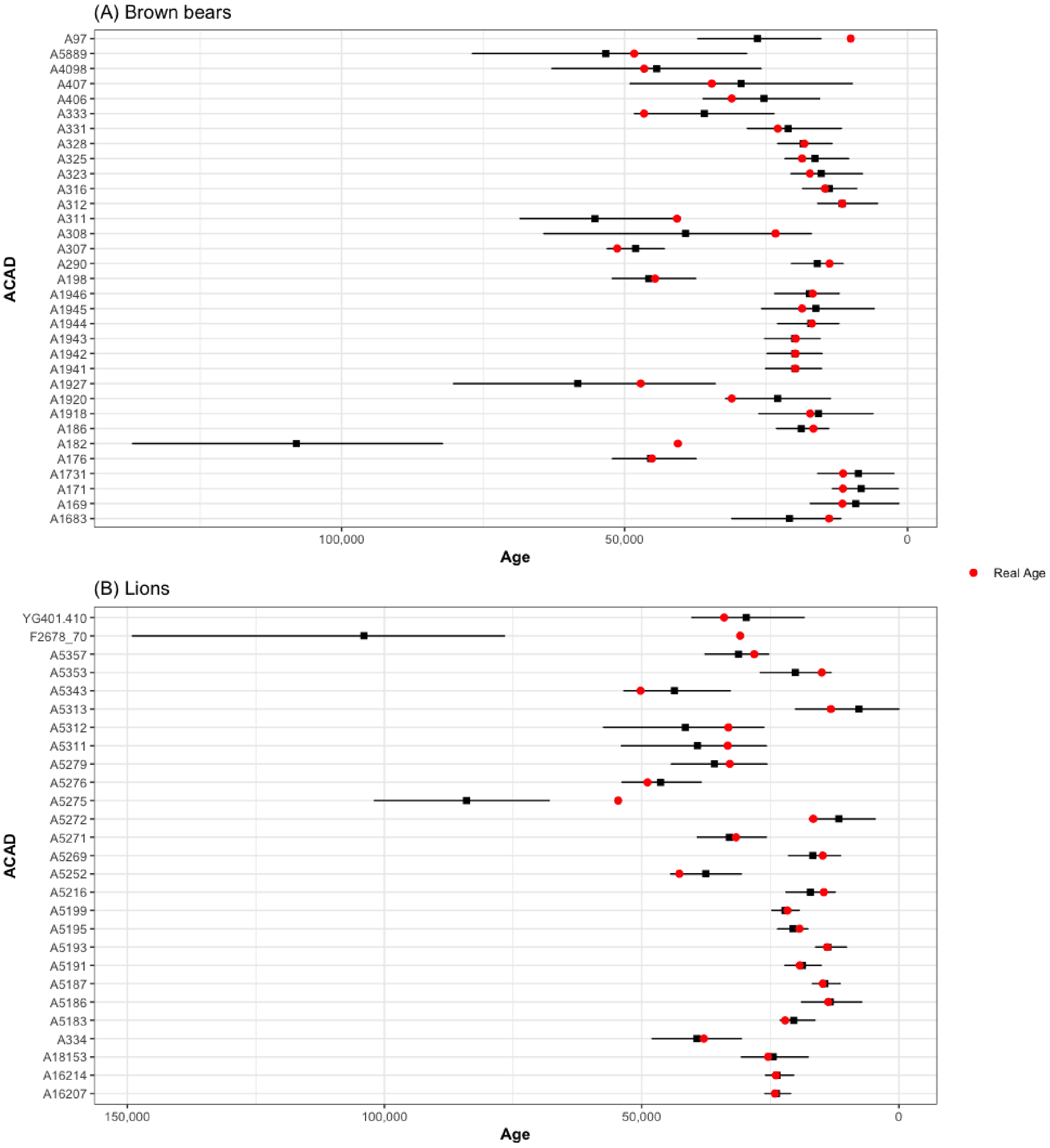


**Figure S1:** Plots of median estimated ages from leave-one-out cross-validation in BEAST2 for (A) Brown bears and (B) Lions. Error-bars represent 95% higher posterior density (HPD). The real age of the specimen is within the 95% HPD of each estimate for all but 2 of each of the brown bears and lions. The specimens for which the real age is outside the 95% HPD were still included in subsequent analyses as they fall in under sampled regions of the tree.

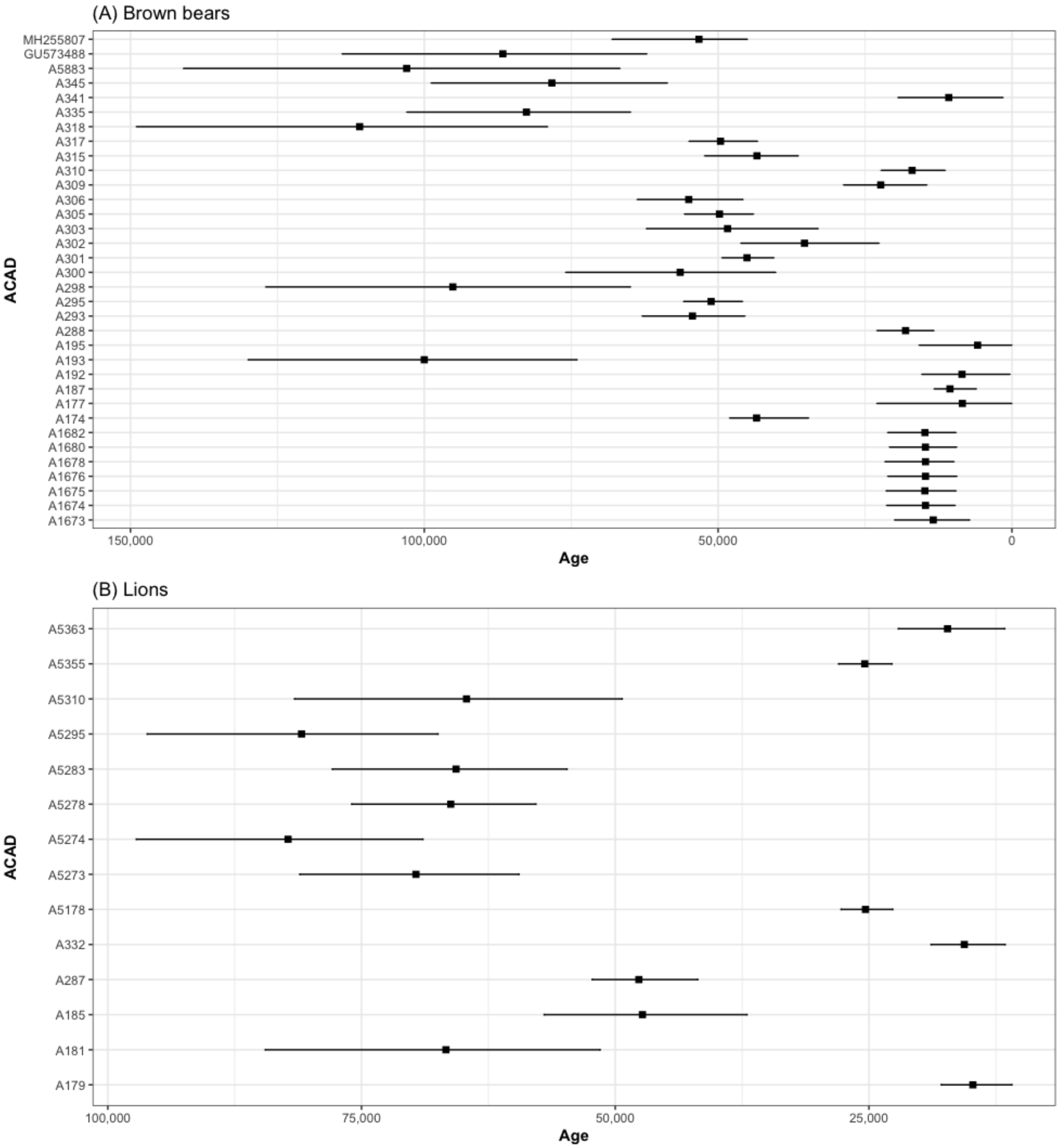


**Figure S2:** Estimated ages from BEAST2 of specimens with no associated date or infinite radiocarbon dates. Error bars represent 95% higher posterior densities.

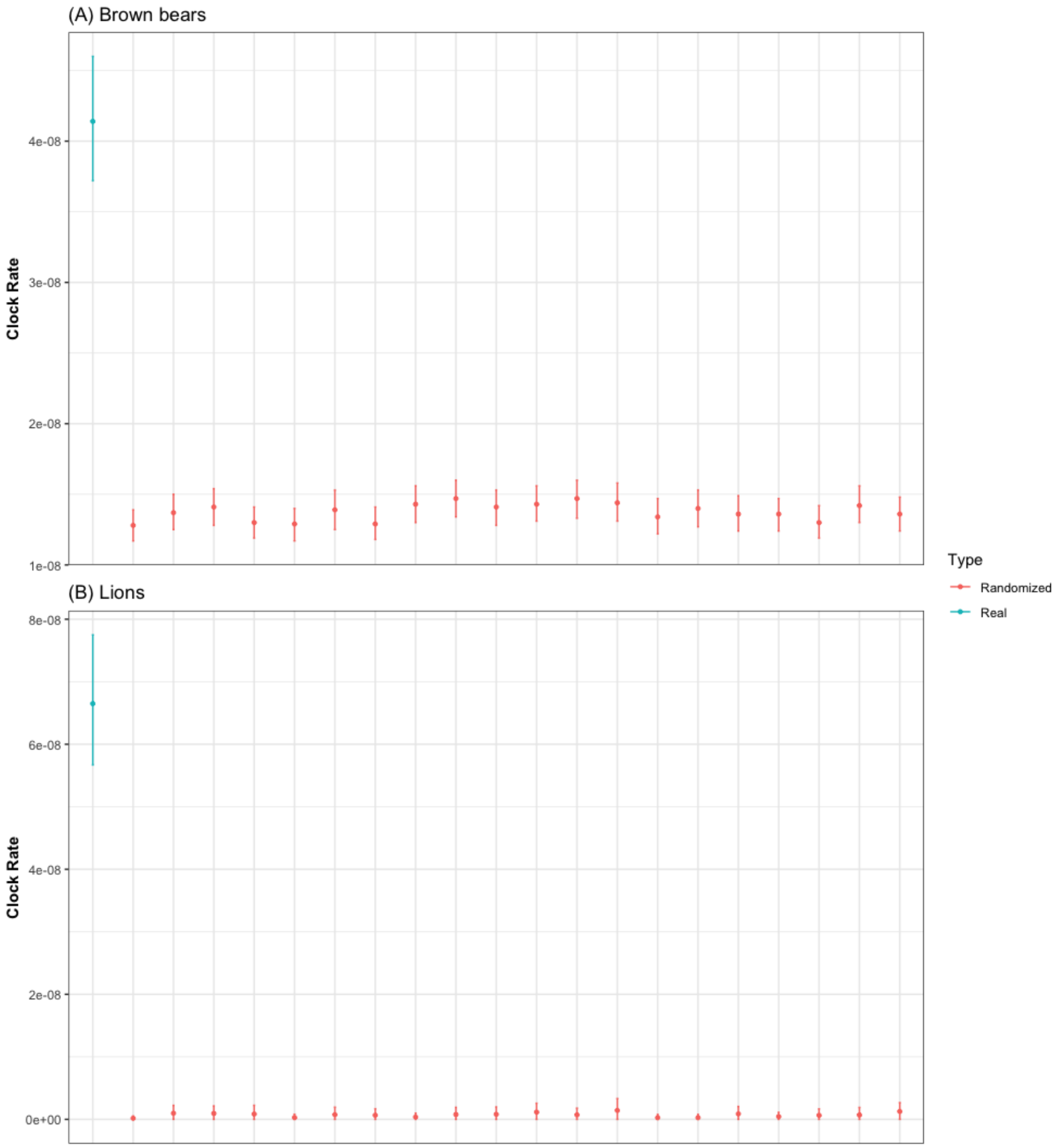


**Figure S3:** Comparison of mean clock rate estimations with 95% higher posterior intervals from BEAST2 for the real data and the 20 date-randomized datasets from the date-randomization test (DRT).

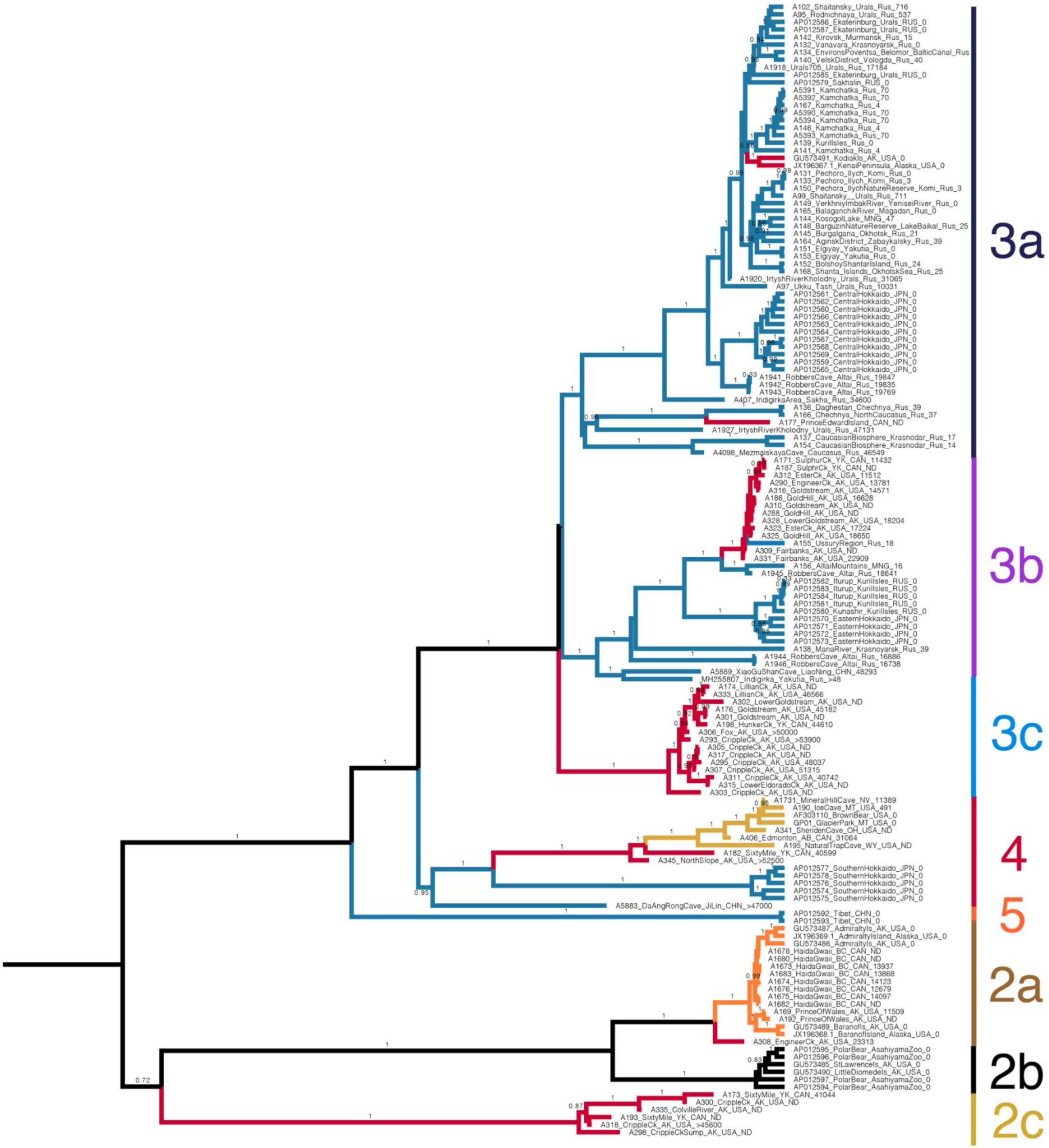


**Figure S4:** Bayesian phylogenetic tree inferred from brown bear mitogenomes. Branch labels represent posterior support values above 0.75.

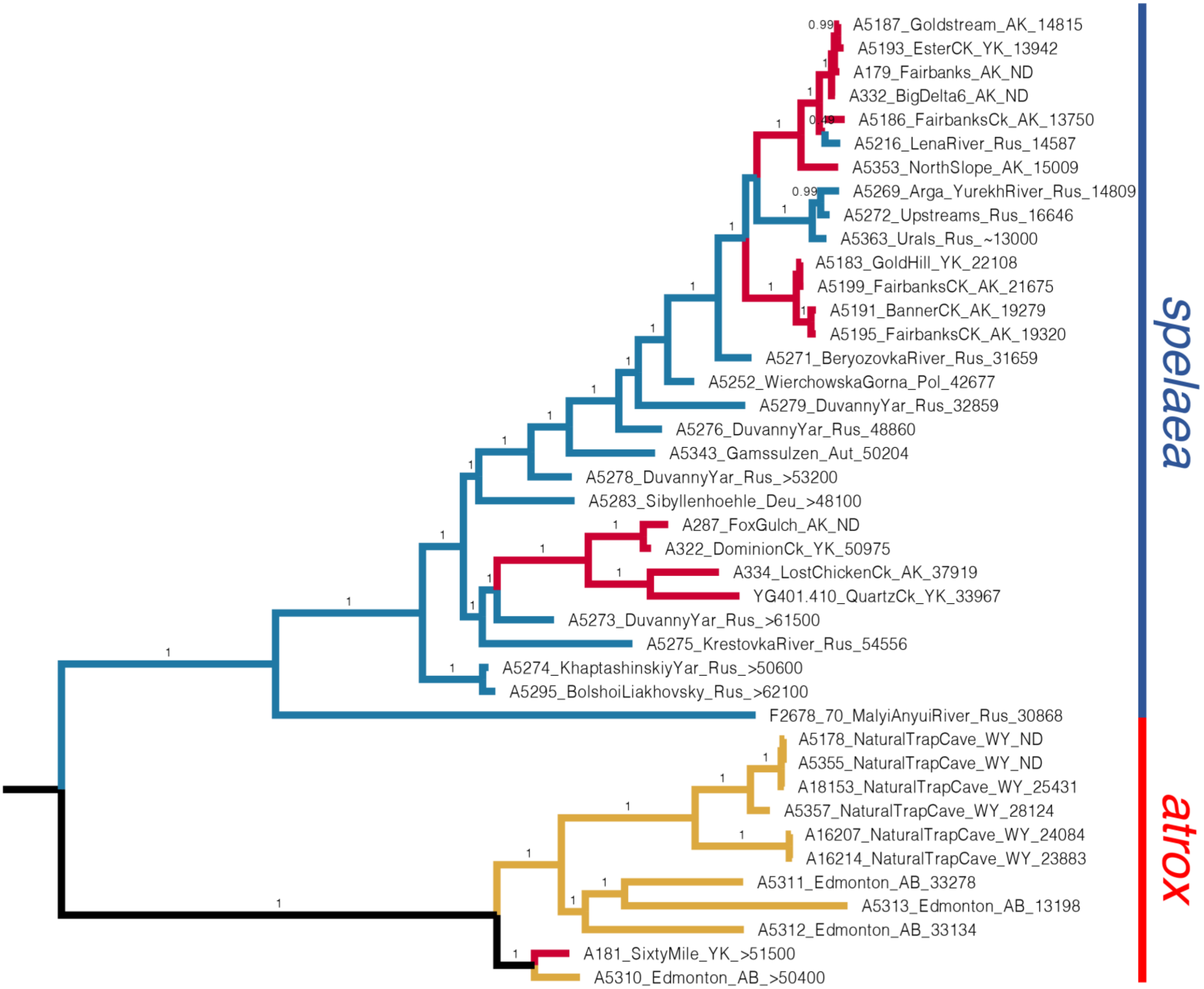


**Figure S5:** Bayesian phylogenetic tree inferred from lion mitogenomes. Branch labels represent posterior support values above 0.75.

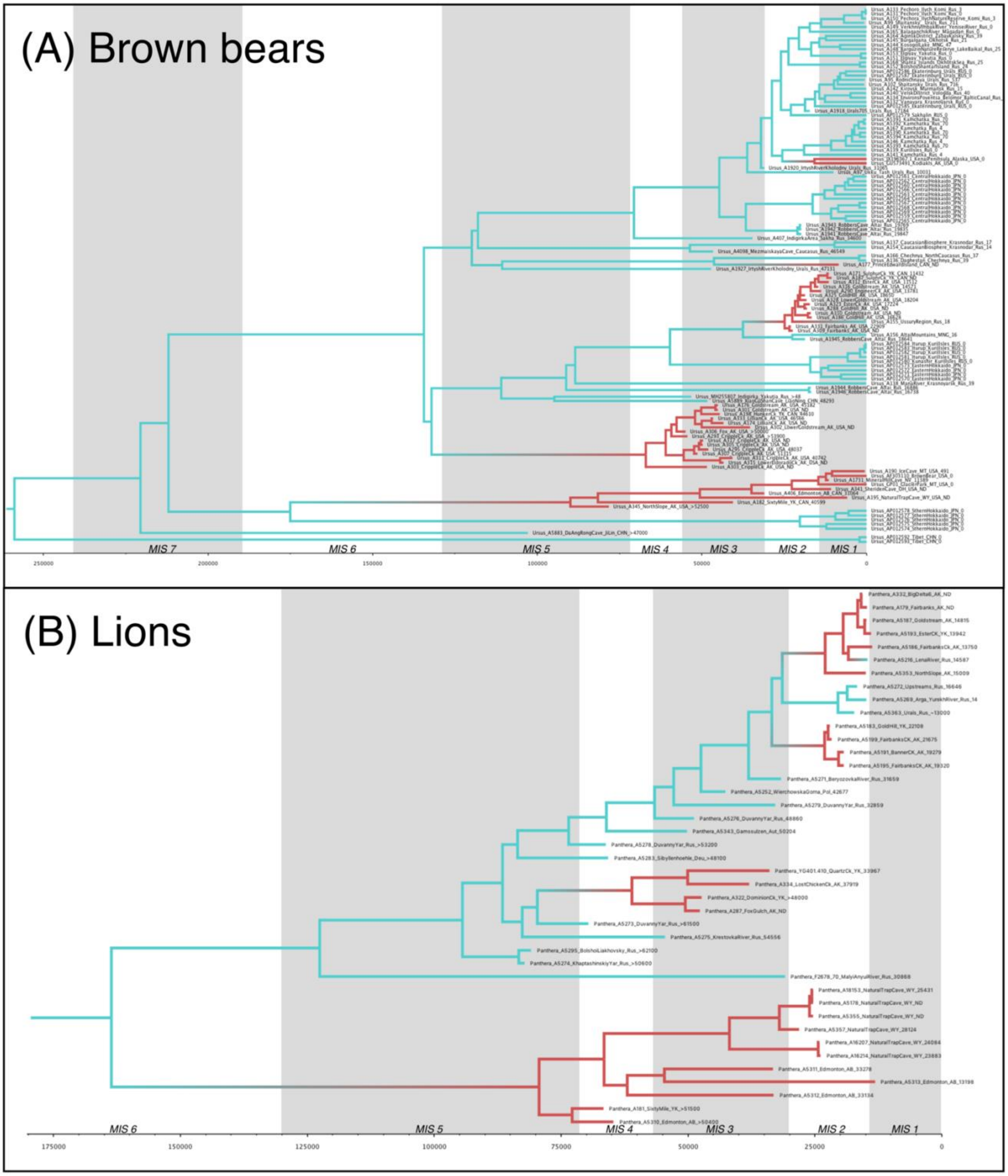


**Supplementary Figure 6:** Bayesian phylogenetic trees inferred from (A) brown bear and (B) lion mitogenomes under joint-tree epoch analyses in BEAST. The grey vertical columns represent odd-numbered MIS stages (interglacials) and white columns even-numbered MIS stages (glacials). Colours of branches correspond to geographic character in the joint-tree epoch analyses, where blue is Eurasia and red North America. Shifts from blue to red and vice versa are inferred migrations across the Bering Land Bridge

**Table S1:** Additional radiocarbon dates used to produce Figure 2.

| Date ID | Species | Museum Accession | Locality | Country | Radiocarbon age | Radiocarbon error | Calibrated median | Calibrated error |
|---|---|---|---|---|---|---|---|---|
| ANUA-38615 | *A.simus* | YT03/134 | Quartz Creek, Yukon | Canada | 26940 | 570 | 31029 | 576 |
| OxA-37428 | *A.simus* | YT03/288 Cat No 129.1 | Hester Creek, Klondike, Dawson, Yukon | Canada | 26800 | 240 | 30950 | 157 |
| OxA-9259 | *A.simus* | CMN 49874 | Dawson area, Hester Creek Loc.57, Yukon | Canada | 26720 | 270 | 30899 | 193 |
| Wk20235 | *A.simus* | CMN 37957 | Dawson area Loc. 45, Eldorado Creek, Yukon | Canada | 22417 | 452 | 26713 | 436 |
| OxA-37426 | *A.simus* | A-1828 | Goldstream, Alaska | USA | 20900 | 120 | 25233 | 188 |
| TO-2699 | *A.simus* | CMN 42388 | Sixtymile, Yukon | Canada | 44240 | 930 | 47621 | 984 |
| OxA-37425 | *A.simus* | A-203-2808 | Cripple Creek, Alaska | USA | 44600 | 2000 | 47763 | 1347 |
| I-11037 | *A.simus* | CMN 37577 | Lower Hunker Creek, Yukon | Canada | 29600 | 1200 | 33744 | 1384 |
| Wk20236 | *A.simus* | AMNH A-'Alaska' Bx 35 | Alaska | USA | 25264 | 650 | 29450 | 695 |
| AA-17511 | *A.simus* | NA | Fairbanks, Alaska | USA | 20524 | 180 | 24727 | 262 |
| AA-17512 | *A.simus* | NA | Fairbanks, Alaska | USA | 25496 | 224 | 29632 | 330 |
| AA-17513 | *A.simus* | NA | Upper Cleary Creek Fairbanks area, Alaska | USA | 27511 | 279 | 31355 | 258 |
| AA-17514 | *A.simus* | NA | Fairbanks, Alaska | USA | 39565 | 1126 | 43536 | 964 |
| AA-17515 | *A.simus* | NA | Birch Creek , Alaska | USA | 34974 | 652 | 39576 | 714 |
| CAMS-58092 | *A.simus* | T99-033 | Titaluk River, Alaska | USA | 42600 | 2200 | 46368 | 1767 |
| TO-2539 | *A.simus* | ROM:VP 43646 | Ikpikpuk River, Alaska | USA | 27190 | 280 | 31158 | 183 |
| TO-2696 | *A.simus* | CMN 7438 | Gold Run Creek (Dawson Loc. 31), Yukon | Canada | 26040 | 270 | 30284 | 336 |
| TO-3707 | *A.simus* | CMN 50367 | Hunker Creek (Dawson Loc. 37), Yukon | Canada | 24850 | 150 | 28884 | 184 |
| OxA-9261 | *U. arctos* | FAM 30771 | Lower Goldstream, Alaska | USA | 20080 | 160 | 24149 | 200 |
| OxA-9709 | *U. arctos* | FAM 95659 | Goldstream, Alaska | USA | 13415 | 70 | 16141 | 115 |
| OxA-9799 | *U. arctos* | FAM 95598 | Cripple Creek, Alaska | USA | 12320 | 90 | 14347 | 216 |
| OxA-9800 | *U. arctos* | FAM 95653 | Engineer Creek, Alaska | USA | 9535 | 75 | 10883 | 144 |
| OxA-9801 | *U. arctos* | FAM 95599 | Goldstream, Alaska | USA | 14310 | 100 | 17430 | 153 |
| OxA-9828 | *U. arctos* | FAM 95628 | Lower Goldstream, Alaska | USA | 12310 | 65 | 14285 | 174 |
| OxA-9829 | *U. arctos* | FAM 95681 | Fairbanks Creek, Alaska | USA | 20820 | 120 | 25119 | 208 |

| **Date ID** | **Species** | **Museum Accession** | **Locality** | **Country** | **Radiocarbon age** | **Radiocarbon error** | **Calibrated median** | **Calibrated error** |
|---|---|---|---|---|---|---|---|---|
| OxA-9830 | *U. arctos* | FAM 95632 | Rosie Creek, Alaska | USA | 14810 | 80 | 18019 | 113 |
| CAMS-131346 | *P. spelaea* | FAM 69126 | Fairbanks, Alaska | USA | 16650 | 110 | 20094 | 157 |
| CAMS-131347 | *P. spelaea* | FAM 69173 | Fairbanks, Alaska | USA | 14050 | 80 | 17077 | 152 |
| CAMS-131348 | *P. spelaea* | FAM 69053 | Fairbanks, Alaska | USA | 13040 | 70 | 15616 | 142 |
| CAMS-131349 | *P. spelaea* | FAM 69078 | Fairbanks, Alaska | USA | 18270 | 130 | 22134 | 159 |
| CAMS-131350 | *P. spelaea* | FAM 69080 | Fairbanks, Alaska | USA | 12990 | 70 | 15536 | 136 |
| CAMS-131361 | *P. spelaea* | AMNH 69142 | Fairbanks, Alaska | USA | 18590 | 130 | 22462 | 153 |
| CAMS-131362 | *P. spelaea* | AMNH 69172 | Fairbanks, Alaska | USA | 17140 | 110 | 20678 | 152 |
| CAMS-18421 | *P. spelaea* | NA | Porcupine River, Yukon | Canada | 39300 | 1000 | 43286 | 830 |
| OxA-10085 | *P. spelaea* | FAM 69158 | Cripple Creek Sump, Alaska | USA | 53900 | 2300 | 54706 | 3036 |
| AA-48280 | *P. spelaea* | IK01-409 | Ikpikpuk River, Alaska | USA | 12930 | 130 | 15466 | 201 |
| Beta-117142 | *P. spelaea* | IK97-1001 | Ikpikpuk River, Alaska | USA | 35710 | 1180 | 40335 | 1154 |
| Beta-286419 | *P. spelaea* | IK06-18 | Ikpikpuk River, Alaska | USA | 33260 | 230 | 37514 | 434 |
| Beta-331881 | *P. spelaea* | MAY12-24 | Maybe Creek, Alaska | USA | 15990 | 60 | 19301 | 109 |
| Beta-339277 | *P. spelaea* | TIT12-07 | Titaluk River, Alaska | USA | 30520 | 180 | 34483 | 185 |
| CAMS-131360 | *P. spelaea* | AMNH 69140 | Fairbanks, Alaska | USA | 20970 | 180 | 25303 | 243 |
| CAMS-53909 | *P. spelaea* | IK98-278 | Ikpikpuk River, Alaska | USA | 11290 | 50 | 13143 | 51 |
| CAMS-53910 | *P. spelaea* | IK98-436 | Ikpikpuk River, Alaska | USA | 40900 | 1140 | 44558 | 1085 |
| SI-456 | *P. spelaea* | NA | Upper Ester Creek, Alaska | USA | 22680 | 300 | 26965 | 323 |
| TO-7743 | *P. spelaea* | NA | Thistle Creek, Yukon | Canada | 32750 | 370 | 36838 | 552 |

**Table S2:** Information on read data downloaded from EMBL-EBI.

| Sample | SRA number/ EBI run accession number | Age | Location | Species | Mitochondrial Clade | Reference |
|---|---|---|---|---|---|---|
| GP01 | SRR935602, SRR935609, SRR935616, SRR935617, SRR941811, SRR941814 | Modern | Glacier National Park, Montana, USA | *Ursus arctos* | Clade 4 | Liu et al. 2014 |
| F2678/70 | SRR3591801, SRR3591802 | 28,690 ± 130, OZQ292 | Malyi Anyui river, Chukotka, Russia | *Panthera spelaea* | *Spelaea* | Barnett et al. 2016 |
| YG 401.410 | SRR3630971 | 29,860 ± 210, UCIAMS-143525 | Quartz Creek, Yukon, Canada | *Panthera spelaea* | *Spelaea* | Barnett et al. 2016 |

**Table S3:** Information on sequences downloaded from GenBank included as part of the brown bear dataset.

| GenBank code | Age | Location | Species | Mitochondrial Clade | Reference |
|---|---|---|---|---|---|
| AF303110.1 | Modern | Continental USA | *Ursus arctos* | 4 | Delisle & Strobeck 2002 |
| AP012559.1 | Modern | Central Hokkaido | *Ursus arctos* | 3a | Hirata et al. 2013 |
| AP012560.1 | Modern | Central Hokkaido | *Ursus arctos* | 3a | Hirata et al. 2013 |
| AP012561.1 | Modern | Central Hokkaido | *Ursus arctos* | 3a | Hirata et al. 2013 |
| AP012562.1 | Modern | Central Hokkaido | *Ursus arctos* | 3a | Hirata et al. 2013 |
| AP012563.1 | Modern | Central Hokkaido | *Ursus arctos* | 3a | Hirata et al. 2013 |
| AP012564.1 | Modern | Central Hokkaido | *Ursus arctos* | 3a | Hirata et al. 2013 |
| AP012565.1 | Modern | Central Hokkaido | *Ursus arctos* | 3a | Hirata et al. 2013 |
| AP012566.1 | Modern | Central Hokkaido | *Ursus arctos* | 3a | Hirata et al. 2013 |
| AP012567.1 | Modern | Central Hokkaido | *Ursus arctos* | 3a | Hirata et al. 2013 |
| AP012568.1 | Modern | Central Hokkaido | *Ursus arctos* | 3a | Hirata et al. 2013 |
| AP012569.1 | Modern | Central Hokkaido | *Ursus arctos* | 3a | Hirata et al. 2013 |
| AP012570.1 | Modern | Eastern Hokkaido | *Ursus arctos* | 3b | Hirata et al. 2013 |
| AP012571.1 | Modern | Eastern Hokkaido | *Ursus arctos* | 3b | Hirata et al. 2013 |
| AP012572.1 | Modern | Eastern Hokkaido | *Ursus arctos* | 3b | Hirata et al. 2013 |
| AP012573 | Modern | Eastern Hokkaido | *Ursus arctos* | 3b | Hirata et al. 2013 |
| AP012574 | Modern | Southern Hokkaido | *Ursus arctos* | 4 | Hirata et al. 2013 |
| AP012575 | Modern | Southern Hokkaido | *Ursus arctos* | 4 | Hirata et al. 2013 |
| AP012576 | Modern | Southern Hokkaido | *Ursus arctos* | 4 | Hirata et al. 2013 |
| AP012577 | Modern | Southern Hokkaido | *Ursus arctos* | 4 | Hirata et al. 2013 |
| AP012578 | Modern | Southern Hokkaido | *Ursus arctos* | 4 | Hirata et al. 2013 |
| AP012579 | Modern | Sakhalin | *Ursus arctos* | 3a | Hirata et al. 2013 |
| AP012580.1 | Modern | Kunashiri Island | *Ursus arctos* | 3b | Hirata et al. 2013 |
| AP012581.1 | Modern | Etorofu Island | *Ursus arctos* | 3b | Hirata et al. 2013 |
| AP012582.1 | Modern | Etorofu Island | *Ursus arctos* | 3b | Hirata et al. 2013 |
| AP012583.1 | Modern | Etorofu Island | *Ursus arctos* | 3b | Hirata et al. 2013 |
| AP012584.1 | Modern | Etorofu Island | *Ursus arctos* | 3b | Hirata et al. 2013 |
| AP012585 | Modern | Ekaterinburg | *Ursus arctos* | 3a | Hirata et al. 2013 |
| AP012586 | Modern | Ekaterinburg | *Ursus arctos* | 3a | Hirata et al. 2013 |
| AP012587 | Modern | Ekaterinburg | *Ursus arctos* | 3a | Hirata et al. 2013 |
| AP012592.1 | Modern | Tibet | *Ursus arctos* | 5 | Hirata et al. 2013 |
| AP012593.1 | Modern | Tibet | *Ursus arctos* | 5 | Hirata et al. 2013 |
| AP012594.1 | Modern | Asahikawa Municipal Asahiyama Zoo, Japan | *Ursus maritimus* | 2b | Hirata et al. 2013 |
| AP012595.1 | Modern | Asahikawa Municipal Asahiyama Zoo, Japan | *Ursus maritimus* | 2b | Hirata et al. 2013 |

| GenBank code | Age | Location | Species | Mitochondrial Clade | Reference |
|---|---|---|---|---|---|
| AP012596.1 | Modern | Asahikawa Municipal Asahiyama Zoo, Japan | *Ursus maritimus* | 2b | Hirata et al. 2013 |
| AP012597.1 | Modern | Asahikawa Municipal Asahiyama Zoo, Japan | *Ursus maritimus* | 2b | Hirata et al. 2013 |
| GU573485.1 | Modern | St Lawrence Island, Alaska, USA | *Ursus maritimus* | 2b | Lindqvist et al. 2010 |
| GU573486.1 | Modern | Admiralty Island, Alaska, USA | *Ursus arctos* | 2a | Lindqvist et al. 2010 |
| GU573487.1 | Modern | Admiralty Island, Alaska, USA | *Ursus arctos* | 2a | Lindqvist et al. 2010 |
| GU573489.1 | Modern | Baranof Island, Alaksa, USA | *Ursus arctos* | 2a | Lindqvist et al. 2010 |
| GU573490.2 | Modern | Little Diomede Island, Alaska, USA | *Ursus maritimus* | 2b | Lindqvist et al. 2010 |
| GU573491.1 | Modern | Kodiak Island | *Ursus arctos* | 2a | Lindqvist et al. 2010 |
| JX196367.1 | Modern | Kenai Peninsula, Alaska, USA | *Ursus arctos* | 3a | Miller et al. 2012 |
| JX196368.1 | Modern | Baranof Island, Alaksa, USA | *Ursus arctos* | 2a | Miller et al. 2012 |
| JX196369.1 | Modern | Admiralty Island, Alaska, USA | *Ursus arctos* | 2a | Miller et al. 2012 |
| MH255807.1 | >48000 | Indigirka river basin, Uyandina river, Yakutia, Russia | *Ursus arctos* | 3b | Rey-Iglesia et al. 2019 |

**Table S4:** Best partition scheme according to BIC for BEAST analyses

| Brown Bears | | | |
|---|---|---|---|
| Partition | Best Model | #sites | Positions |
| 1 | TRN+I | 4227 | Second codon position of ND6, tRNAs, and rRNAs |
| 2 | HKY+I | 3603 | First codon position of ATP6, ATP8, CO1, CO2, CO3, CYTB, ND1, ND2, ND3, ND4, ND4L, ND5, ND6 |
| 3 | HKY+I | 3603 | Second codon position of ATP6, ATP8, CO1, CO2, CO3, CYTB, ND1, ND2, ND3, ND4, ND4L, ND5, ND6 |
| 4 | TRN+G | 3955 | First and third codon position ND6, third codon position of ATP6, ATP8, CO1, CO2, CO3, CYTB, ND1, ND2, ND3, ND4, ND4L, ND5, ND6 |
| 5 | TRN+I+G | 1063 | Non Coding |
| **Lions** | | | |
| Partition | Best Model | #sites | Positions |
| 1 | HKY | 4229 | Second codon position of ND6, tRNAs, and rRNAs |
| 2 | HKY+I | 3602 | First codon position of ATP6, ATP8, CO1, CO2, CO3, CYTB, ND1, ND2, ND3, ND4, ND4L, ND5, ND6 |
| 3 | HKY+I | 3602 | Second codon position of ATP6, ATP8, CO1, CO2, CO3, CYTB, ND1, ND2, ND3, ND4, ND4L, ND5, ND6 |
| 4 | TRN | 3954 | First and third codon position ND6, third codon position of ATP6, ATP8, CO1, CO2, CO3, CYTB, ND1, ND2, ND3, ND4, ND4L, ND5, ND6 |
| 5 | TRN+I+G | 1357 | Non Coding |

**Dataset S1:** Information on brown bear bone and tooth samples analyzed. New radiocarbon dates are highlighted in red.

See Excel File

**Dataset S2:** Information on lion bone and tooth samples analyzed. New radiocarbon dates are highlighted in red.

See Excel File
